# PSGL-1 blockade delays relapse to BRAF/MEK inhibition in cutaneous melanoma

**DOI:** 10.64898/2026.07.02.736105

**Authors:** OS El Naggar, BN Ha, ML Rakoto, L Cort, A Amirfallah, EA Haglund, P Urquiza, HAF Hetrick, LM Bradley, EJ Hartsough, JL Hope, G Romano

## Abstract

Advanced BRAF-mutant cutaneous melanoma can be treated with targeted therapy when immune checkpoint inhibitors (ICIs) fail or are not a feasible option. Nevertheless, most patients do not achieve a durable response, highlighting the critical need for therapeutic partners that enhance the long-term efficacy of targeted therapy. Transcriptomic analysis of a BRAF-mutant melanoma model of acquired resistance identified P-selectin glycoprotein ligand-1 (PSGL-1) as a top-upregulated immune mediator upon resistance acquisition. PSGL-1 is a key regulator of CD8^+^ T cell exhaustion and differentiation, and its inhibition has been shown to enhance T cell function across multiple disease models. Based on these observations, we hypothesized that combined targeting of BRAF/MEK and PSGL-1 would improve anti-tumor responses.

Here, we demonstrate that dual inhibition of BRAF/MEK and PSGL-1 elicits durable tumor control in a preclinical model of PD-1-refractory cutaneous melanoma. Single-cell RNA sequencing of the tumor microenvironment reveals robust reprogramming of intratumoral CD8^+^ T cells toward a less terminally differentiated, memory-like phenotype following combined BRAF/MEK and PSGL-1 targeting. Consistent with these findings, CD8^+^ T cells in the tumor-draining lymph nodes of PSGL-1^-/-^ mice exhibit enhanced functionality and a less differentiated state of exhaustion when compared with wild-type mice. To extend these observations to a translationally relevant setting, we further show that antibody-mediated blockade of PSGL-1, in combination with BRAF/MEK inhibition, yields superior anti-tumor activity compared with either monotherapy. Collectively, these findings identify PSGL-1 as a promising therapeutic target to enhance the durability of targeted therapy and provide a strong rationale for future clinical evaluation.

## Introduction

BRAF-mutant cutaneous melanoma is an aggressive skin malignancy that frequently becomes resistant to the FDA-approved therapies BRAF/MEK-targeted inhibitors and combined anti-PD-1/anti-CTLA-4 immune checkpoint inhibition (ICI). While ICI is the current standard-of-care for patients with advanced melanoma, targeted therapy (TT) remains valuable as a bridge to immunotherapy and when ICI fails [1]. BRAF/MEK-targeted therapies in cutaneous melanoma induce rapid tumor regression and exert profound T cell infiltration and expression of T cell-specific tumor antigens, thereby promoting a “hot” tumor microenvironment (TME) [2]. However, this enhanced immune activation is transient, and resistance to targeted therapy is associated with the restoration of a suppressive, or “cold,” TME [3, 4]. To leverage increased tumor-specific T cell activation and promote more durable anti-tumor immunity, combinatorial approaches with ICI and BRAF/MEK inhibition have been assessed clinically. However, attempts to combine MAPK-targeted therapy and ICI have resulted in systemic toxicity (most notably when combined with anti-CTLA-4 blockade), and so-called “triplet therapy” has focused on patients with advanced disease for which ICI and TT had already failed [5]. These clinical setbacks support the exploration of novel companion targets for TT to provide effective and durable therapeutic options to patients with BRAF-altered, treatment-refractory melanoma.

P-selectin glycoprotein ligand-1 (PSGL-1) is a glycoprotein that is highly expressed on immune cells. Our previous studies demonstrated that PSGL-1 is a potent immune checkpoint upstream of PD-1 and a critical regulator of TCR signaling and T cell differentiation [6, 7]. While agonistic signaling through PSGL-1 promotes exhaustion in CD8^+^ T cells, genetic deficiency or therapeutic blockade of PSGL-1 signaling restrains terminal differentiation and enhances proliferation, infiltration, and functional capacity. Critically, removal of intrinsic PSGL-1 signaling in CD8^+^ T cells promotes increased retention of the transcription factor TCF-1 and decreased TOX expression, profiles which are associated with increased stemness [7].

CD8^+^ T cells are the primary effector cell type responsible for the direct killing of tumor cells [7]. In the TME, chronic antigen exposure drives an exhausted phenotype in CD8^+^ T cells (T_EX_), which is characterized by loss of effector function. T_EX_ can arise from precursor T cells (T_PEX_) that differentiate into various early, intermediate, and terminally differentiated T cells (T_EX-term_) [8]. However, the mediators of durable response in the TME are memory-like CD8^+^ T cells [9]. These subsets are driven by distinct transcriptional programs that determine T cell function and fate. Defining how BRAF/MEK-targeted therapy reshapes CD8^+^ T cell transcriptional states within the TME will provide insight into the mechanisms governing effector function, exhaustion, and durable anti-tumor immunity [7].

In this study, we show that combined BRAF/MEK inhibition and PSGL-1 targeting induce durable anti-tumor responses in a preclinical model of PD-1–refractory cutaneous melanoma. Both genetic ablation and antibody-mediated blockade of PSGL-1 enhance the efficacy of targeted therapy and prolong relapse-free survival. Molecular analysis reveals that combination treatment reshapes intratumoral CD8^+^ T cell transcriptional programs toward a memory-like state, while enriching polyfunctional CD8^+^ T cells in tumor-draining lymph nodes, collectively supporting durable tumor control following treatment cessation.

## Methods

### Cell Culture

Yumm1.5 (Braf^V600E/wt^ Pten^−/−^Cdkn2^−/−^) cells were cultured in complete media consisting of DMEM (Dulbecco’s Modified Eagle Medium)/Hams F-12 50/50 Mix [+] L-glutamine (Corning #10-092-CV) supplemented with 10% fetal bovine serum (FBS, Avantor #89510-186) and 1X Antibiotic-Antimycotic Solution (Corning #30-004-CI). Cells were maintained at 37°C, 5% CO_2_. For in vivo studies, Yumm1.5 cells were passaged 2-3 times prior to injection. Cells were routinely monitored and tested for mycoplasma contamination using a Mycoplasma PCR Detection Kit (Abcam #ab289834).

### Mice

C57BL/6 (CD90.2^+^) mice and PSGL-1^-/-^ (CD90.1^+^) mice (B6.Cg-*Selplg*^*tm1Fur*^/J) were either bred in house (C57BL/6, PSGL-1^-/-^) or purchased (C57BL/6) from Jackson Laboratories. All mice were housed in the animal facility at Drexel University College of Medicine. All animal experiments were performed under the Institutional Animal Care and Use Committee (IACUC) at Drexel University College of Medicine-approved protocols (LA-23-019 and LA-24-129). For long-term in vivo studies, 7–17-week-old male mice were used. For flow cytometry, single-cell RNASeq, and antibody experiments, 6–9-week-old male mice were used.

### Tumor Studies

For tumor injections, cells were washed once with Phosphate-Buffered Saline (1X PBS), 1X Without Calcium and Magnesium (Corning #21-040-CV), and incubated in 0.25% Trypsin/ 2.21mM EDTA 1X HBSS without sodium bicarbonate, calcium, and magnesium (Corning #25-053-CI) for approximately 2-3 minutes before quenching with complete media. Following detachment, cells were washed once with 1X PBS and counted. Cells were centrifuged for 5 min at 300 x g at room temperature (RT) and resuspended at the appropriate volume in a 1:1 solution of PBS and Matrigel Basement Membrane Matrix, LDEV-free (Corning #354234). Unless otherwise described, 5x10^5^ Yumm1.5 cells were intradermally injected in the right hind flank following shaving in a total volume of 100 µL per animal. All tumors were allowed 14-16 days to develop (average volume 50-80mm^3^) until the onset of targeted therapy. For all in vivo tumor studies, mice were weighed 3 times per week, and tumors were measured on the indicated days. Tumors were measured using calipers, and tumor volume (in mm^3^) was calculated using the following formula: Volume = p/6 x (L x W x H), where L = length, W = width, and H = height or depth of the tumor. BRAF/MEK inhibitor was administered as follows: Open standard diet with 15 kcal% Fat (Research Diets, Inc.) synthesized with the following drugs: 200 ppm PLX-4720 (TargetMol Cat. # T2473) and 7ppm PD0325901 (Mirdametinib, TargetMol Cat. # T6189). For antibody studies, anti-PD-1 mAb (clone RMP1-14, BioXCell #BE0146) or rat IgG2a isotype control (clone 2A3, BioXCell #BE0089) were administered by intraperitoneal injections once per week for 4 weeks (200µg/mouse). A novel anti-mouse PSGL-1 monoclonal antibody (mAb) (see below) was administered intraperitoneally (I.P.) as described (200µg/mouse).

### Antibody Generation and Validation

Novel anti-PSGL-1 antibodies were generated by immunizing mice with purified recombinant mouse PSGL-1 protein to generate Fab libraries, which were then selected based on binding to PSGL-1. The selected Fabs were then formatted and produced as IgG mAbs and screened for high binding affinity and specificity. To validate anti-PSGL1 mAb specificity, spleens were harvested from one C57BL/6 and one PSGL-1^-/-^ mouse and passed through a 45 µm cell strainer to generate a single-cell suspension. Splenocytes were treated with 1X red blood cell lysis buffer (Biolegend, Cat. #422401) for 1 minute. Following washing, single-cell suspensions of splenocytes were incubated with varying concentrations of the anti-mouse PSGL-1 antibody for 20 minutes at 4°C. Cells were washed and incubated with BV421-conjugated anti-mouse IgG (BV421, Jackson ImmunoResearch, Cat. #715-675-151, RRID #AB_2651117). Cells were then incubated with a cocktail of surface stain antibodies: Alexa Fluor® 488 anti-mouse CD4 (Biolegend, Cat. #100532, RRID #AB_493373), PE/Cyanine7 anti-mouse CD8a (Biolegend, Cat. #100722, RRID #AB_312761), and Brilliant Violet 605™ anti-mouse CD19 (Biolegend, Cat. #115540, RRID #AB_2563067). Samples were washed and resuspended in 1% formaldehyde in PBS and analyzed on a BD LSRFortessa™ flow cytometer.

### Flow Cytometry

At indicated time points, mice were euthanized, and tissues were harvested (spleen, tumor, and tumor-draining and non-draining inguinal lymph nodes) and placed in RPMI 1640 (Corning Cat. #10-040-CV) + 5% FBS on ice. Lymph nodes and spleens were passed through a 40µm cell strainer (Fisherbrand), and splenocytes were treated with 1X red blood cell lysis buffer (Biolegend, Cat. #422401) for 1 minute. Tumors were digested using the gentleMACS™ tissue dissociator (Miltenyi Biotec) in 2.5 mL of a tumor digestion cocktail (250 U/mL Collagenase CLSAFA, 20 U/mL Collagenase CLSAFC, 20 U/mL Deoxyribonuclease I in RPMI 1640). Tumors were then incubated at 37°C with gentle rocking for an additional 30 minutes. The tumor digestion cocktail was quenched with RPMI 1640 + 5% FBS. All cells were washed and counted using a Countess cell counter (Thermo Fisher) and plated in sterile V-bottom 96-well dishes for stimulation. Samples containing fewer than 1x10^6^ cells were pooled for stimulation and analysis. Cells were stimulated overnight (approximately 16 hours) with 10 U/mL human IL-2, 1 μg/mL anti-CD3 (BioXCell Cat. #BE0001-1), and 0.5 µg/mL anti-mouse CD28 (BioXCell Cat. #BE0015-1) in complete T cell media (RPMI 1640 supplemented with 55 µM 2-mercaptoethanol and Brefeldin A). The next day, cells were washed once with FACSWash (1X HBSS + 5% FBS), incubated with anti-mouse CD16/CD32 (BioXCell Cat. #BE0307) for 15 minutes at 4°C and incubated with a cocktail of surface stain antibodies (Table S1). Cells were suspended in Foxp3 intracellular fixation buffer and then permeabilized (eBioscience, ThermoFisher Cat. #00-5523-00). Following permeabilization, cells were incubated with a cocktail of intracellular antibodies (**Table S1**). The cells were then washed and resuspended in 1% formaldehyde in PBS and filtered through a 40µm cell strainer prior to analysis. All analysis for tumor experiments was conducted on the BD FACSymphony™ A5 SE analyzer and analyzed using FlowJo V10.10 (BD Biosciences).

### Single-cell RNA Sequencing

Tumors were harvested and digested as described above. Dissociated tumors were subjected to dead cell removal (Miltenyi Biotec Cat. #130-090-101) according to the manufacturer’s protocol. Samples were subsequently fixed using the Parse Biosciences fixation kit and stored at ^−^80 °C for 48 hours per the manufacturer’s instructions. A total of 53,520 cells across 12 samples were loaded into the Parse Biosciences Evercode WT v3 barcoding workflow. Samples were barcoded and processed into 8 sublibraries according to the manufacturer’s protocol and sequenced at Parse Biosciences. The quality and concentration of the sublibraries were measured using a TapeStation 2100 Bioanalyzer (Agilent).

Raw count matrices were generated using the Parse Biosciences analysis pipeline and imported into R (version 4.4.2) for downstream analysis. Single-cell analysis was performed using Seurat (version 5.5.0). Sparse count matrices were used to generate a Seurat object retaining genes detected in at least three cells and cells containing at least 200 detected genes. Quality control filtering excluded cells with fewer than 200 or greater than 8,000 detected genes and cells with mitochondrial transcript content exceeding 20%. Putative doublets were identified using scDblFinder (version 1.20.2) implemented through the SingleCellExperiment framework following initial normalization and dimensionality reduction. Cells classified as doublets were removed prior to downstream analyses. Following doublet removal, data were re-normalized using Seurat’s “LogNormalize” method with a scale factor of 10,000. Highly variable genes (n = 4,000) were identified using the variance-stabilizing transformation (“vst”) method. Data were scaled and subjected to principal component analysis (PCA) using the top 30 principal components. To correct for inter-sample technical variation while preserving biological heterogeneity, datasets were integrated using Harmony (version 2.0.2) based on sample identity.

Dimensionality reduction was performed using Uniform Manifold Approximation and Projection (UMAP) on Harmony-corrected embeddings. Graph-based clustering was performed using Seurat’s FindNeighbors() and FindClusters() functions using the first 20 Harmony dimensions and a clustering resolution of 1.0, resulting in transcriptionally distinct cell populations. Cluster-specific marker genes were identified using the FindAllMarkers() function employing a Wilcoxon rank-sum test with a minimum expression threshold of 25% of cells and a log fold-change threshold >0.25. Cell populations were annotated manually based on canonical lineage markers and transcriptional profiles, including T cell subsets, tumor-associated macrophages, dendritic cells, neutrophils, natural killer cells, cancer-associated fibroblasts, B cells, mast cells, and tumor cells (**Table S2**). Low-quality or ambiguous clusters were excluded from downstream analyses.

Samples were grouped into four experimental conditions (WT_Veh, WT_BM, KO_Veh, and KO_BM) based on metadata annotations. Cell-type proportions were quantified across groups using normalized frequencies. Differential gene expression analyses within selected populations were performed using Seurat’s FindMarkers() function. For CD8^+^ T cell trajectory analyses, CD8^+^ T cell subsets were isolated and converted into SingleCellExperiment objects. Trajectory inference was performed using Slingshot (version 2.14.0) on UMAP embeddings with biologically defined progenitor-like CD8 T cell populations specified as the trajectory root state (**Table S3**). Multiple differentiation trajectories were identified, including terminally exhausted, proliferative exhausted, and memory-like lineages (**Table S4**). Pseudotime values were projected back onto Seurat objects for downstream visualization and analysis. Genes associated with lineage progression were identified using Spearman correlation analyses between gene expression and lineage-specific pseudotime values.

Visualization and statistical analyses were performed in R using Seurat, ggplot2, dplyr, viridis, slingshot, SingleCellExperiment, tidyverse, and related packages. Visualizations included UMAP projections, violin plots, feature plots, trajectory overlays, and cell composition analyses.

### Statistics

Statistical analyses were performed in GraphPad Prism, and unless otherwise specified, data are presented as mean ± SEM. Data distributions were analyzed (Kolmogorov-Smirnov, Shapiro-Wilk) before applying parametric tests. Multi-group comparisons used ANOVA or Kruskal-Wallis, followed by Tukey’s or Dunn’s post-hoc tests. For in vivo tumor studies, two-way repeated-measures ANOVA (or a mixed-effects model if missing values) was performed, with multiple comparisons followed by the two-stage step-up method of Benjamin, Krieger, and Yekutieli (FDR 0.05). Kaplan-Meier curves were analyzed using Benjamini-Hochberg for multiple comparisons (FDR 0.05). Absolute cell counts, ratios, and frequencies from flow cytometry data were analyzed using ordinary one-way ANOVA with multiple comparisons followed by Tukey’s tests.

### Data availability

Bulk transcriptomic (GSE79972) and scRNAseq datasets are accessible through GEO (accession number pending). All other data are available within the figures and Supplementary Materials or from the corresponding authors upon request.

## Results

### Genetic deletion of PSGL-1 impairs relapse to BRAF/MEK targeted therapy in a mouse model of melanoma

To identify novel companion targets for use in combination with targeted therapies, we analyzed our previously published transcriptomic dataset [10] of a mouse model of BRAF^V600E^ melanoma (iBIP; inducible-Braf^V600E^, Ink/Arf^-/-^, Pten^-/-^) [11]. The iBIP model features a doxycycline-inducible Braf^V600E^ oncogene. Activation of the Braf^V600E^ oncogene induces melanomagenesis, and its extinction by removing doxycycline causes rapid immune activation and significant tumor regression. Following tumor regression, minimal residual disease (MRD) is established, causing an immunologically “cold” tumor microenvironment (TME). iBIP tumors exhibit low T cell infiltration and are unresponsive to both PD-1 and CTLA-4 ICIs. Our analysis of time-course transcriptomic data showed that P-selectin glycoprotein ligand-1 (PSGL-1, encoded by the *Selplg* gene) is the most upregulated immune checkpoint following BRAF extinction (**Fig. 1A, Fig. S1A**). In addition, PSGL-1 transcripts remain elevated through MRD (**Fig. 1A**).

**Figure 1.**
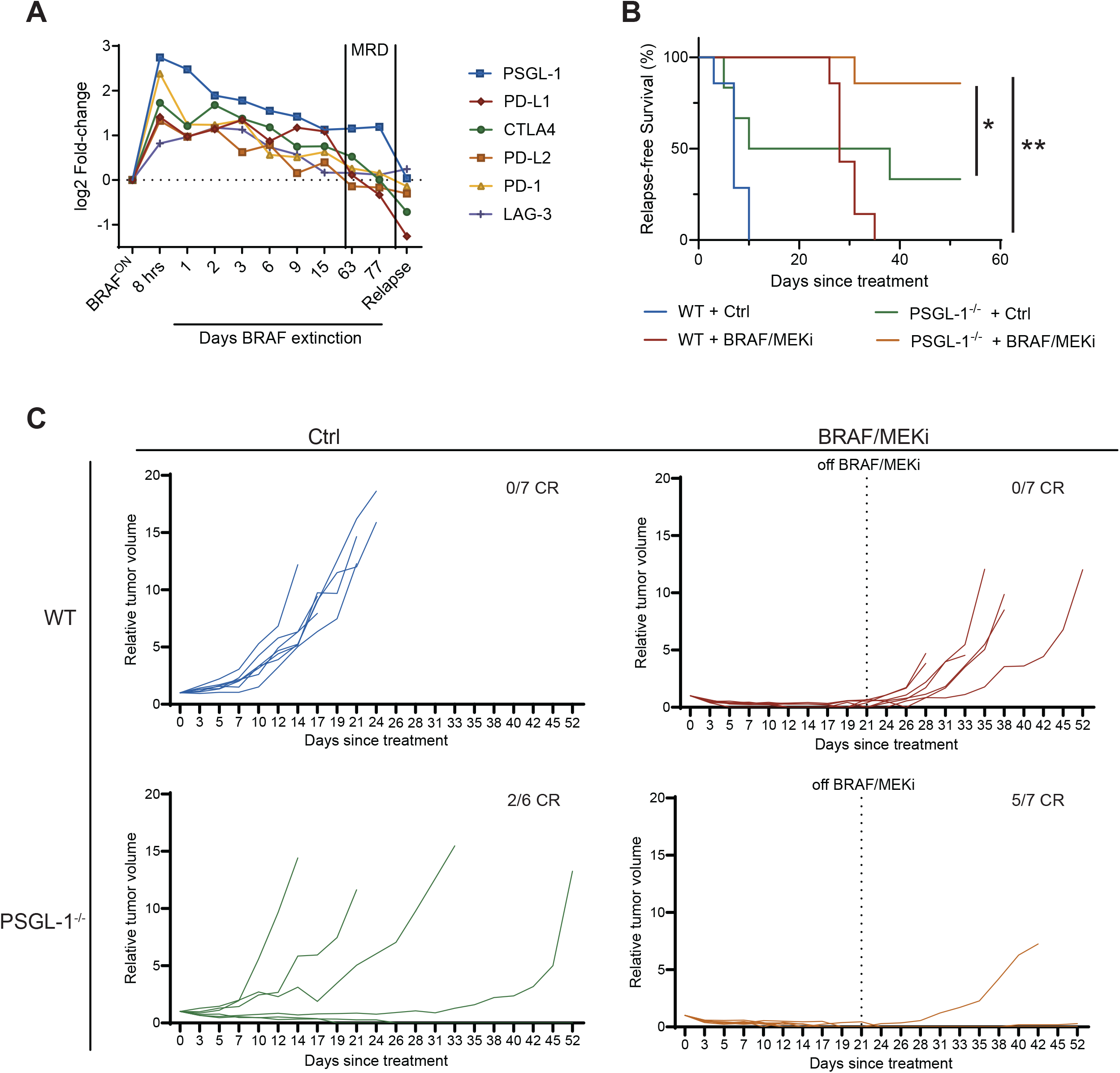
Relapse following BRAF/MEK inhibition is ablated in a genetic model of PSGL-1 blockade in Yumm1.5 melanoma. **(A)** Top upregulated immune checkpoints after genetic BRAFV600E extinction (days) in the iBIP model. Data are median centered and reported as log2 fold change. MRD, minimal residual disease. **(B)** Yumm1.5 cells (5x10^5^ cells) were intradermally injected in the flank of male wild-type or PSGL-1^-/-^ mice (n= 6 or 7 per group). When average tumor volume reached ∼60mm^3^, BRAF/MEKi or Vehicle control chow were administered. BRAF/MEKi was discontinued after 3 weeks and tumors were monitored for relapse. Kaplan-Meier analysis of time to relapse of Yumm1.5 tumors following administration of BRAF/MEKi (time to relapse was recorded as days until tumors reached a volume of 100mm^3^ following BRAF/MEKi). **(C)** Relative tumor volumes of wildtype or PSGL1^-/-^ mice +/- BRAF/MEKi. Individual datapoints are shown for each mouse. Tumor volume values were normalized to day 0 on BRAF/MEKi. Dotted line indicates removal of BRAF/MEKi or Vehicle control chow. Number of mice per group that reached complete response (CR) (defined by lack of palpable tumor) are displayed in the top right of each graph. * for P ≤ 0.05, ** for P ≤ 0.01, ns (not significant) for P ≥ 0.05.

Given the sustained expression of PSGL-1 in MRD following BRAF^V600E^ extinction, we hypothesized that combined targeting of PSGL-1 and BRAF/MEK would yield a durable anti-tumor response in murine melanoma. To answer this question, we used a genetic model of PSGL-1 blockade – a full-body knockout of the *Selplg* gene (PSGL-1^-/-^). We and others have shown that PSGL-1^-/-^ mice exhibit significant tumor control with respect to the syngeneic, PD-1-refractory mouse melanoma tumor line, Yumm1.5 (**Fig. S1B**) [12]. We administered Yumm1.5 cells intradermally into the flank of male wildtype (WT) or PSGL-1 knockout (PSGL-1^-/-^) mice and administered BRAF/MEKi (open standard diet + 7ppm PD0325901 + 200ppm PLX-4720) or vehicle control chow (Ctrl) for 21 days (beginning 14 days after Yumm1.5 inoculation). In wild-type mice, Yumm1.5 tumors rapidly rebounded following discontinuation of BRAF/MEKi (**Fig. 1C**). In contrast, we observed a complete response in 5 out of 7 PSGL-1^-/-^ mice treated with BRAF/MEK-targeted therapy (**Fig. 1C, Table S5**) – complete response was measured as the absence of a palpable tumor. Furthermore, the observed tumor control with BRAF/MEKi administration in PSGL-1 knockout mice was durable for at least 45 days (**Fig. S1C**) following discontinuation of TT, with mice showing no gross evidence of tumor upon dissection. When analyzed as relapse-free survival (measured as days until tumor volume reached 100mm^3^ after treatment), combined BRAF/MEKi and PSGL-1 deficiency showed a profound response in 6/7 mice compared to PSGL-1^-/-^ (2/6) or BRAF/MEKi (0/7) alone (**Fig. 1B**). Taken together, these data support the hypothesis that combined BRAF/MEK and PSGL-1 targeting improves treatment durability in a model of PD-1- and BRAF/MEKi-resistant melanoma.

### scRNA-seq reveals robust reprogramming of the T cell compartment in the tumor microenvironment under concomitant BRAF/MEK and PSGL-1 targeting

To define immune dynamics in melanoma following BRAF/MEK inhibition and/or PSGL-1 deletion, we performed single-cell RNA sequencing of Yumm1.5 tumors from WT and PSGL-1^-/-^ mice 72 hours after targeted therapy (n = 3 biological replicates/group, **Fig S2A-B**). Unsupervised clustering of integrated single-cell transcriptomes identified 27 transcriptionally distinct clusters (**Fig S2C**). These clusters were subsequently annotated into major tumor, immune, and stromal cell populations based on the expression of canonical marker genes (**Fig. 2A, Fig. S2D, Table S2**). The T cell compartment was the most abundant across all treatment groups (**Fig. 2B**). To further resolve T cell heterogeneity, the T cell compartment was subsetted and reclustered, identifying ten subclusters corresponding to distinct differentiation states of CD4^+^ and CD8^+^ T cells based on canonical marker expression and transcriptional annotations (**Fig. 2C, Table S3, Fig. S3A-C**). We defined several subsets related to exhaustion (T_EX_), including early and late precursor exhausted T cells (T_PEX-early_, T_PEX-late_), which can sustain response and self-renew under chronic antigen exposure and differentiate into other T_EX_ subsets (**Fig. 2C**). Further differentiated intermediate/effector-like T_EX_ states were also identified (T_EX-eff_), along with more specific subsets that are enriched in transcriptional programs associated with proliferation, stress response, and interferon response (T_EX-prolif_, T_EX-int-stress_ and T_EX-int-IFN_) (**Fig. 2C**). Further differentiated were the CD8-T_EX-term_, representing terminally exhausted or dysfunctional T cells, which sustain limited proliferative capacity and diminished effector function (**Fig. 2C**). Across groups, CD8-T_EX-term_ were the most abundant T cell subset but were at the highest proportion in the BRAF/MEKi-treated WT group (**Fig. S3D)**. Trajectory inference analysis of the CD8^+^ T cell compartment identified three major differentiation programs corresponding to terminally exhausted, proliferative, and memory-like states (**Fig. 2D, Table S4, Fig. S4A-C**). To characterize the transcriptional dynamics underlying these trajectories, we examined the expression of canonical T cell state markers across pseudotime (**Fig. 2E, Fig. S4D**). As expected, *Mki67*, which encodes the proliferation marker Ki-67, was strongly induced along the proliferative trajectory while remaining comparatively low in terminal and memory-like lineages. In contrast, *Tcf7*, encoding the stemness-associated transcription factor TCF-1, was broadly expressed early in pseudotime across all trajectories but was sustained within the memory-like lineage at later differentiation states. Expression of *Pdcd1* and *Tox*, two hallmark regulators of T cell exhaustion, was enriched along terminal and proliferative differentiation trajectories, consistent with progressive acquisition of dysfunctional or exhausted transcriptional programs [13]. Importantly, memory-like CD8^+^ T cells were most abundant in the PSGL-1^-/-^ mice treated with BRAF/MEKi compared to any other group (**Fig. S3D**). Quantification of lineage occupancy revealed a significant enrichment of the memory-like trajectory specifically in the BRAF/MEKi-treated PSGL-1^-/-^ group, whereas other treatment groups remained biased toward terminally exhausted or proliferative trajectories (**Fig. 2F**). These findings suggest that dual PSGL-1 ablation and BRAF/MEKi promote CD8^+^ T cell differentiation toward a memory-associated transcriptional state rather than terminal exhaustion.

**Figure 2.**
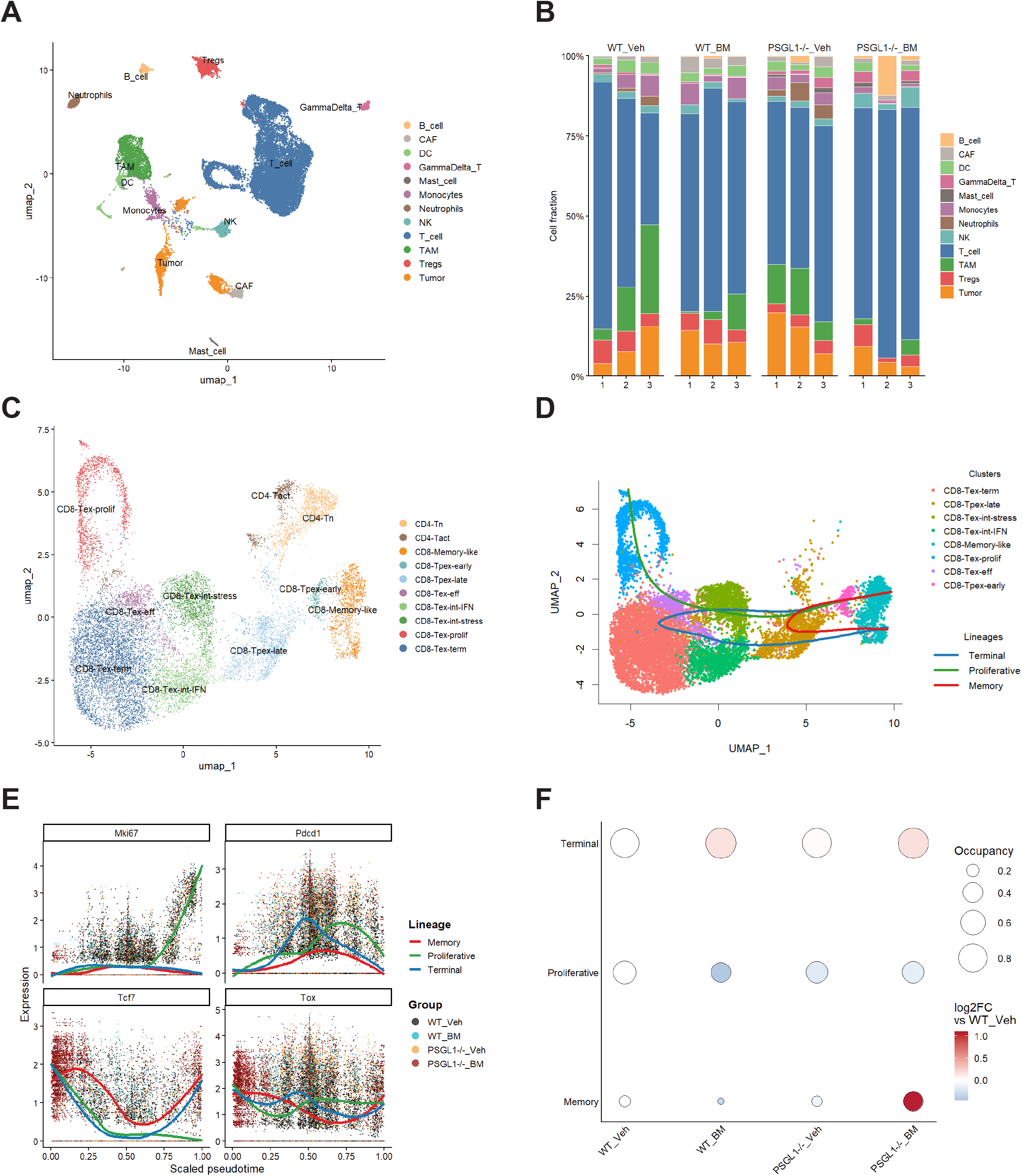
scRNAseq reveals remodeling of the T cell compartment following PSGL-1 deletion and BRAF/MEK inhibition. **(A)** UMAP visualization of integrated scRNA-seq data from Yumm1.5 tumors harvested 72 h after treatment from WT and PSGL-1^-/-^ mice receiving vehicle or BRAF/MEK inhibitor (BM) therapy (n=3 mice per group). Unsupervised clustering identified major tumor, immune, and stromal cell populations annotated using canonical marker genes. **(B)** Relative abundance of annotated cell populations across experimental groups and biological replicates. Each bar represents a single tumor sample, and colors correspond to the cell populations shown in (A).**(C)** UMAP projection of tumor-infiltrating T cells following subclustering analysis. Unsupervised clustering identified transcriptionally distinct T-cell states, including naïve, activated, interferon-responsive, proliferative, exhausted, terminally exhausted, and memory-like populations. **(D)** Slingshot trajectory inference overlaid on the CD8+ T-cell UMAP, revealing three major differentiation lineages corresponding to terminal, proliferative, and memory-like cell states. Curves indicate inferred lineage trajectories across pseudotime. **(E)** Expression dynamics of representative lineage-associated genes (*Mki67, Pdcd1, Tcf7*, and *Tox*) along scaled pseudotime. Individual points represent single cells colored by experimental group, and solid lines indicate lineage-specific smoothed expression trends. **(F)** Summary of Slingshot trajectory analysis across experimental groups. Bubble size denotes lineage occupancy (fraction of CD8^+^ T cells assigned to each lineage), whereas color indicates the relative change in lineage representation compared with WT vehicle-treated controls (log2 fold change).

### PSGL-1 deletion enhances CD8^+^ T cell effector function following BRAF/MEK inhibition in the tumor-draining lymph nodes

Given the robust changes that PSGL-1 deficiency confers to the CD8^+^ T cell compartment in the context of BRAF/MEKi, as well as an established T cell dependency for PSGL-1’s role in the Yumm1.5 model [6], we focused our studies on validating functional changes in the CD8^+^ T cell population in the tumor-draining lymph node (tdLN) in response to combination therapy. We sought to characterize early immune reprogramming in the tdLN following TT initiation, given that the lymph node is the key site for the activation of naïve T cells prior to migration into the tumor [14, 15]. 48 hours after BRAF/MEKi treatment initiation, flow cytometry was used to profile immunological changes in the tdLN (**Fig. S5A, Fig. S6A-C**). We observed increases in absolute numbers of both CD4^+^ and CD8^+^ T cells in the tdLN of PSGL-1^-/-^ mice compared to WT controls (**Fig. 3A**), regardless of BRAF/MEKi exposure. We also observed a significant increase in the CD8:CD4 T-cell ratio in the tdLN of PSGL-1^-/-^ mice (**Fig. 3B**). In line with previous findings, PSGL-1-deficiency promoted an increase in the frequency of cytokine-producing CD8^+^ T cells compared to wild-type CD8^+^ T cells (**Fig. 3C, D**). In the context of BRAF/MEK inhibition, treatment with BRAF/MEKi alone induced an approximately two-fold increase in the frequency of TNFα single-positive CD8^+^ T cells relative to WT controls (**Fig. 3E**). In contrast, PSGL-1 deficiency preferentially promoted the maintenance of polyfunctional IFNγ^+^TNFα^+^CD8^+^ T cells, independent of BRAF/MEK inhibition (**Fig. 3F**). Notably, combined PSGL-1 loss and BRAF/MEK inhibition preserved elevated frequencies of double-positive polyfunctional CD8+ T cells. Together, these findings suggest that while BRAF/MEK inhibition primarily enhances a TNFα effector response, PSGL-1 ablation sustains a broader polyfunctional CD8^+^ T cell state characterized by concurrent IFNγ and TNFα production.

**Figure 3.**
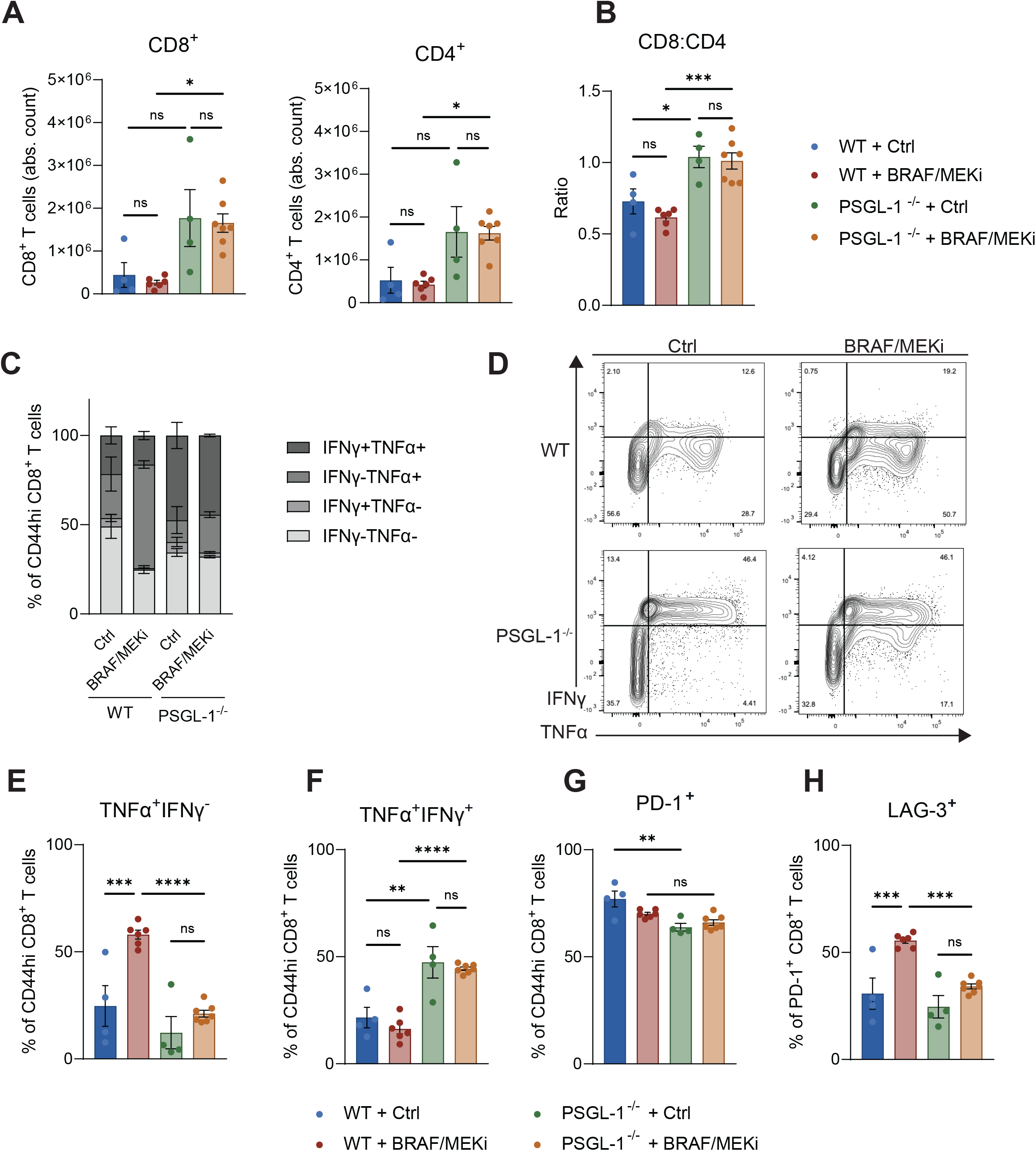
Polyfunctional CD8^+^ T cells are enriched in the tumor-draining lymph node (tdLN) following BRAF/MEK inhibition under PSGL-1 deletion. **(A)** Absolute cell counts of CD4^+^ and CD8^+^ T cells gated out of CD90.1^+^ and CD90.2^+^ cells from the tumor-draining lymph node of wildtype (WT) or PSGL-1^-/-^ mice, 48 hours after BRAF/MEKi administration (n= 6 or 7 mice per group, samples with fewer than 1x10^6^ cells were pooled). Cells were stimulated with anti-CD3 and anti-CD28 mAb in the presence of IL-2 and brefeldin A and analyzed via flow cytometry. **(B)** Ratio of the absolute counts of CD8^+^ to CD4^+^ T cells from (A). **(C)** Frequency of double-positive or -negative or single-positive IFNγ- or TNFα-producing CD8^+^ T cells from tdLN following stimulation. **(D)** Representative flow cytometry contour plots showing one sample from each group from data shown in (C). **(E and F)** Frequency of double positive or TNFα-producing CD8^+^ T cells from each group, from data shown in (C). **(G)** Frequency of PD-1-expressing CD8^+^ T cells in tdLN following stimulation. **(H)** Frequency of LAG-3-expressing cells, gated out of PD-1^+^CD8^+^ T cells in tdLN following stimulation. All data are presented as mean ± SEM. * for P ≤ 0.05, ** for P ≤ 0.01, *** for P ≤ 0.001, and **** for P ≤ 0.0001, ns (not significant) for P ≥ 0.05.

Next, we hypothesized that PSGL-1-mediated alterations in T cell differentiation may be critical in driving enhanced tumor control. In evaluating T cell phenotypes, we observed no difference in the frequency of PD-1^+^ CD8^+^ T cells between Ctrl and BRAF/MEKi-treated WT mice (**Fig. 3G**). However, PD-1 levels were significantly lower in PSGL-1^-/-^ mice than in WT mice, as previously observed [7]. The frequency of LAG-3-expressing PD-1^+^ CD8^+^ T cells was increased in WT mice treated with BRAF/MEKi (consistent with tumor data in **Fig. 1A**), but not so in PSGL-1^-/-^ groups (**Fig. 3H**). No significant changes in TIM-3- or CX3CR1-expressing PD-1^+^ CD8^+^ T cells were observed (**Fig. S6D-E**). These findings indicate that loss of PSGL-1 signaling promotes an early, more functionally active CD8^+^ T cell state in the tDLN, characterized by enhanced effector capacity.

### Antibody-based PSGL-1 blockade delays the relapse to BRAF/MEK targeted therapy

Given the profound anti-tumor response we observed when combining genetic PSGL-1 deficiency with BRAF/MEK-targeted therapy, we sought to evaluate the translational potential of this novel combinatorial treatment strategy. Using a novel function-blocking anti-mouse PSGL-1 IgG1 mAb, we tested sequential inhibition of PSGL-1 followed by inhibition of BRAF/MEK (**Fig. S7A**). As before, we injected Yumm1.5 cells intradermally into the flanks of male C57BL/6 mice. When tumors had reached an average volume of 50 mm^3^, we began antibody treatment. To compare to a standard-of-care immune checkpoint inhibitor, we treated mice with 6 cycles of once weekly anti-PD-1 (clone RMP1-14, 200 µg/mouse, I.P.), and control groups were treated with rat IgG2a isotype control (clone 2A3, 200 µg/mouse, I.P.) following the same schedule (**Fig. 4A**). In groups treated with anti-PSGL-1, mice were treated 3 times in an antibody “loading” phase every other day following injection with Yumm1.5 cells. BRAF/MEKi (open standard diet + 7ppm PD0325901 + 200ppm PLX-4720) or vehicle control chow (Ctrl) was administered 14 days after Yumm1.5 injection, and animals were treated with a once weekly “maintenance” dose of anti-PSGL-1, anti-PD-1, or isotype control. After 21 days of either targeted therapy or maintenance immunotherapy, one additional loading phase of anti-PSGL-1 was administered to groups receiving anti-PSGL-1. Following discontinuation of all therapies, mice were monitored for relapse, and body weight was measured until humane endpoint (**Fig. 4A** and **Fig S7B**). As expected, neither anti-PD-1 nor anti-PSGL-1 monotherapy significantly reduced tumor burden when compared to isotype-treated mice in the long term (**Fig. S7C**). Despite the limited impact of anti-PSGL-1 as a monotherapy in this approach, we observed that combined BRAF/MEKi and anti-PSGL-1 significantly delayed relapse compared with BRAF/MEKi alone (**Fig. 4B** and **Table S6**). Combined BRAF/MEKi and anti-PD-1 therapy also significantly delayed relapse compared with BRAF/MEKi alone. While this anti-PD1 response was not significantly different from the anti-PSGL-1 combination arm in terms of relapse after BRAF/MEKi discontinuation (**Fig. 4B** and **Table S6**), mice receiving anti-PSGL-1 exhibited superior tumor control to anti-PD-1 and isotype controls in the first 7 days of treatment before BRAF/MEKi initiation (**Table S7**). Collectively, these findings support PSGL-1 blockade as a translationally relevant therapeutic strategy that can stall tumor growth upfront and enhance the durability of responses to BRAF/MEK inhibition in melanoma.

**Figure 4.**
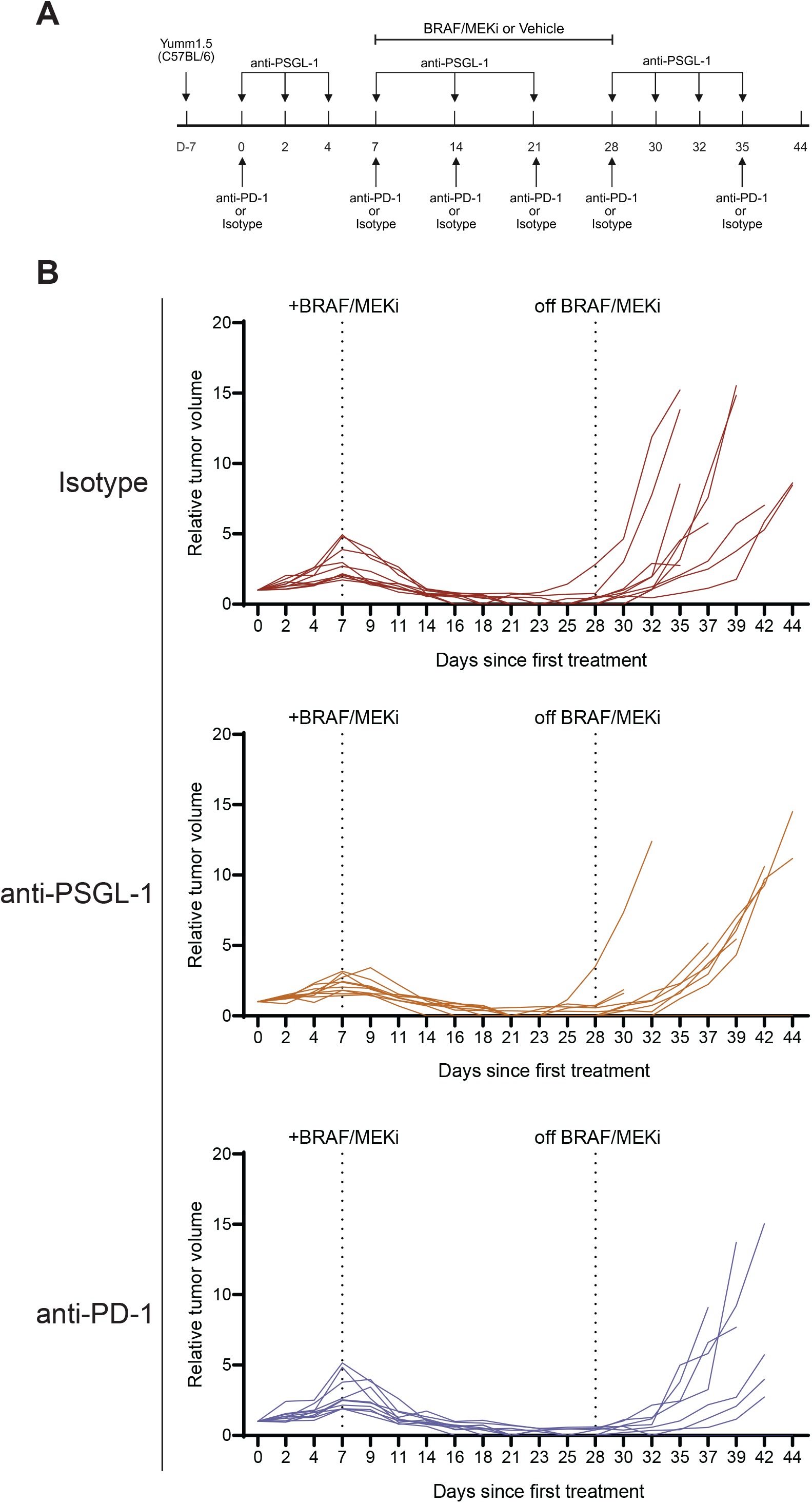
Antibody-based PSGL-1 blockade improves treatment efficacy of BRAF/MEK targeted therapy. **(A)** Experimental timeline of anti-PSGL-1 mAb, anti-PD-1 mAb, or Isotype control in combination with BRAF/MEKi in the Yumm1.5 model. Created in https://BioRender.com. **(B)** Relative tumor volume growth curves of C57BL/6 mice treated with BRAF/MEKi. Yumm1.5 cells (5x10^5^cells) were intradermally injected in the flank of male mice n=10/group). Mice were treated with anti-PD-1, Isotype control, or anti-PSGL-1 as indicated in (A). Tumor volume values were normalized to the first day of antibody treatment (day 7 after tumor injection). Arrows indicate days that mice were treated with anti-PSGL-1. Individual datapoints are shown for each mouse. Dotted line indicate administration or removal of BRAF/MEKi chow.

## Discussion

This study demonstrates, for the first time, the potential of combining PSGL-1 with BRAF/MEKi, leveraging a novel immune checkpoint regulator as a promising tool for melanoma treatment. Considering our previous studies of PSGL-1 as a critical regulator of T cell signaling, our findings position PSGL-1 as a mechanistically distinct and therapeutically complementary immune checkpoint within the current standard of care for melanoma treatment [7]. Unlike PD-1 or CTLA-4, which primarily serve as negative regulators after T cell activation and within the tumor microenvironment, naïve T cells express PSGL-1 and can therefore be targeted earlier during initial T cell activation and early T cell fate differentiation [16]. This distinction is particularly relevant when considering combination with targeted therapies such as BRAF/MEK inhibition, which transiently enhance tumor immunogenicity and T cell infiltration but fail to sustain durable immune control as a monotherapy.

Our previous work in PSGL-1-mediated regulation of T cell responses revealed that PSGL-1 signaling modulates T cell receptor (TCR) signaling strength, resulting in significant changes in T cell metabolism, effector function, and differentiation [7]. Consistent with these findings, our scRNA-sequencing analysis reveals substantial remodeling of the intratumoral T cell compartment following combined BRAF/MEK and PSGL-1 targeting, including the expansion of a memory-like CD8^+^ T cell population. This observation is supported by prior work showing an increased frequency of TCF-1^+^ CD8^+^ T cells in PSGL-1-deficient mice compared with wild-type controls [7]. Importantly, independent studies have also reported upregulation of TCF-1 in CD8^+^ T cells from melanoma patients treated with BRAF/MEK inhibitors [17]. Given the well-established role of TCF-1 in maintaining progenitor-like and memory CD8^+^ T cell states [18], these findings collectively suggest that combined PSGL-1 and BRAF/MEK inhibition promotes the preservation of less differentiated CD8^+^ T cells. Supporting this model, the present study demonstrates that PSGL-1 deletion enhances CD8^+^ T cell function in tumor-draining lymph nodes following BRAF/MEK inhibition, while forestalling differentiation/exhaustion. We therefore propose that PSGL-1 blockade restrains premature T cell differentiation within the tumor-draining lymph node, while BRAF/MEK inhibition increases tumor immunogenicity and antigen availability. Together, these complementary effects create conditions that favor the expansion of memory-like, polyfunctional CD8^+^ T cells that subsequently infiltrate tumors. Furthermore, both PSGL-1 inhibition and BRAF/MEK inhibition increase T cell infiltration into melanoma lesions, as previously demonstrated [3, 7]. The combined enhancement of T cell priming, effector function, and tumor infiltration may therefore underlie the significantly extended durable melanoma control observed with dual targeting.

A key component of our study is the use of a novel PSGL-1-blocking antibody, which serves as proof of concept for the therapeutic potential of inhibiting PSGL-1 signaling to enhance responsiveness to BRAF/MEK-targeted therapy. Of note, we designed our study on the rationale that inhibiting PSGL-1 signaling prior to BRAF/MEKi-driven enhancement of immunogenicity would promote a more robust anti-tumor immune response. Notably, our anti-PSGL-1 antibody did not outperform anti-PD-1 in combination with BRAF/MEKi in terms of the duration of tumor control, but performed better than anti-PD-1 prior to the initiation of targeted therapy. These results suggest that novel, enhanced iterations of the antibody treatments might be needed to more closely match the performance of the knock-out model, and that future studies emphasizing differential impacts on tumor control due to timing, sequence, and dosing may yield critical insights into the underlying mechanisms that support combination success rather than failure.

Potential for toxicity when combining PSGL-1 and BRAF/MEK inhibition is a concern in the clinical setting. Unlike PD-1 or CTLA-4, PSGL-1 is expressed on most immune cells and plays a critical role as a negative regulator, preventing immunopathology under highly inflammatory conditions such as systemic viral infection. However, no adverse events have been observed with PSGL-1 deficiency or blockade across numerous solid tumor models, and no signs of toxicity were detected in this study, even when combined with continuous, sustained BRAF/MEKi administration.

Collectively, this study supports a model in which PSGL-1 functions as a critical upstream early regulator of T cell differentiation and provides a strong rationale for its inclusion in next-generation combinatorial therapy that blocks its function in future clinical trials.

## Funding

G.R. is supported by: WW Smith Charitable Trust - Medical Research Program (#C2507); Department of Defense - Congressionally Directed Medical Research Programs, Melanoma Research Program (ME220048); American Cancer Society (RSG-25-1517461-01-ET); Sidney Kimmel Comprehensive Cancer Center (#00023568 and #901333). J.H. is supported by: Melanoma Research Alliance (#1263917). E.J.H is supported by: National Institutes of Health (R01-CA282901); Sidney Kimmel Comprehensive Cancer Center; Department of Pharmacology & Physiology at Drexel University College of Medicine.

## Conflict of interest statement

The authors declare no conflicts of interest.

## Notes

### Competing Interest Statement

The authors have declared no competing interest.

## References

1. Braden, J., et al., Do BRAF-targeted therapies have a role in the era of immunotherapy? ESMO Open, 2025. 10(7): p. 105314.

2. Frederick, D.T., et al., BRAF inhibition is associated with enhanced melanoma antigen expression and a more favorable tumor microenvironment in patients with metastatic melanoma. Clin Cancer Res, 2013. 19(5): p. 1225–31.

3. Wilmott, J.S., et al., Selective BRAF inhibitors induce marked T-cell infiltration into human metastatic melanoma. Clin Cancer Res, 2012. 18(5): p. 1386–94.

4. Kuske, M., et al., Immunomodulatory effects of BRAF and MEK inhibitors: Implications for Melanoma therapy. Pharmacol Res, 2018. 136: p. 151–159.

5. Albrecht, L.J., et al., Anti-PD-(L)1 plus BRAF/MEK inhibitors (triplet therapy) after failure of immune checkpoint inhibition and targeted therapy in patients with advanced melanoma. Eur J Cancer, 2024. 202: p. 113976.

6. Tinoco, R., et al., PSGL-1 Is an Immune Checkpoint Regulator that Promotes T Cell Exhaustion. Immunity, 2016. 44(6): p. 1470.

7. Hope, J.L., et al., PSGL-1 attenuates early TCR signaling to suppress CD8(+) T cell progenitor differentiation and elicit terminal CD8(+) T cell exhaustion. Cell Rep, 2023. 42(5): p. 112436.

8. Masopust, D., et al., Guidelines for T cell nomenclature. Nat Rev Immunol, 2026. 26(4): p. 298–313.

9. Gebhardt, T., S.L. Park, and I.A. Parish, Stem-like exhausted and memory CD8(+) T cells in cancer. Nat Rev Cancer, 2023. 23(11): p. 780–798.

10. Romano, G., et al., Microparticle-Delivered Cxcl9 Prolongs Braf Inhibitor Efficacy in Melanoma. Cancer Immunol Res, 2023. 11(5): p. 558–569.

11. Kwong, L.N., et al., Co-clinical assessment identifies patterns of BRAF inhibitor resistance in melanoma. J Clin Invest, 2015. 125(4): p. 1459–70.

12. Meeth, K., et al., The YUMM lines: a series of congenic mouse melanoma cell lines with defined genetic alterations. Pigment Cell Melanoma Res, 2016. 29(5): p. 590–7.

13. Wen, S., et al., TCF-1 maintains CD8(+) T cell stemness in tumor microenvironment. J Leukoc Biol, 2021. 110(3): p. 585–590.

14. Dammeijer, F., et al., The PD-1/PD-L1-Checkpoint Restrains T cell Immunity in Tumor-Draining Lymph Nodes. Cancer Cell, 2020. 38(5): p. 685–700 e8.

15. Wei, J., et al., Immune microenvironment of tumor-draining lymph nodes: insights for immunotherapy. Front Immunol, 2025. 16: p. 1562797.

16. Tinoco, R., et al., PSGL-1: A New Player in the Immune Checkpoint Landscape. Trends Immunol, 2017. 38(5): p. 323–335.

17. Peiffer, L., et al., BRAF and MEK inhibition in melanoma patients enables reprogramming of tumor infiltrating lymphocytes. Cancer Immunol Immunother, 2021. 70(6): p. 1635–1647.

18. Chen, Z., et al., TCF-1-Centered Transcriptional Network Drives an Effector versus Exhausted CD8 T Cell-Fate Decision. Immunity, 2019. 51(5): p. 840–855 e5.

