## Supplementary Figures for "PSGL-1 blockade delays relapse to BRAF/MEK inhibition in cutaneous melanoma"

**Figure S1**

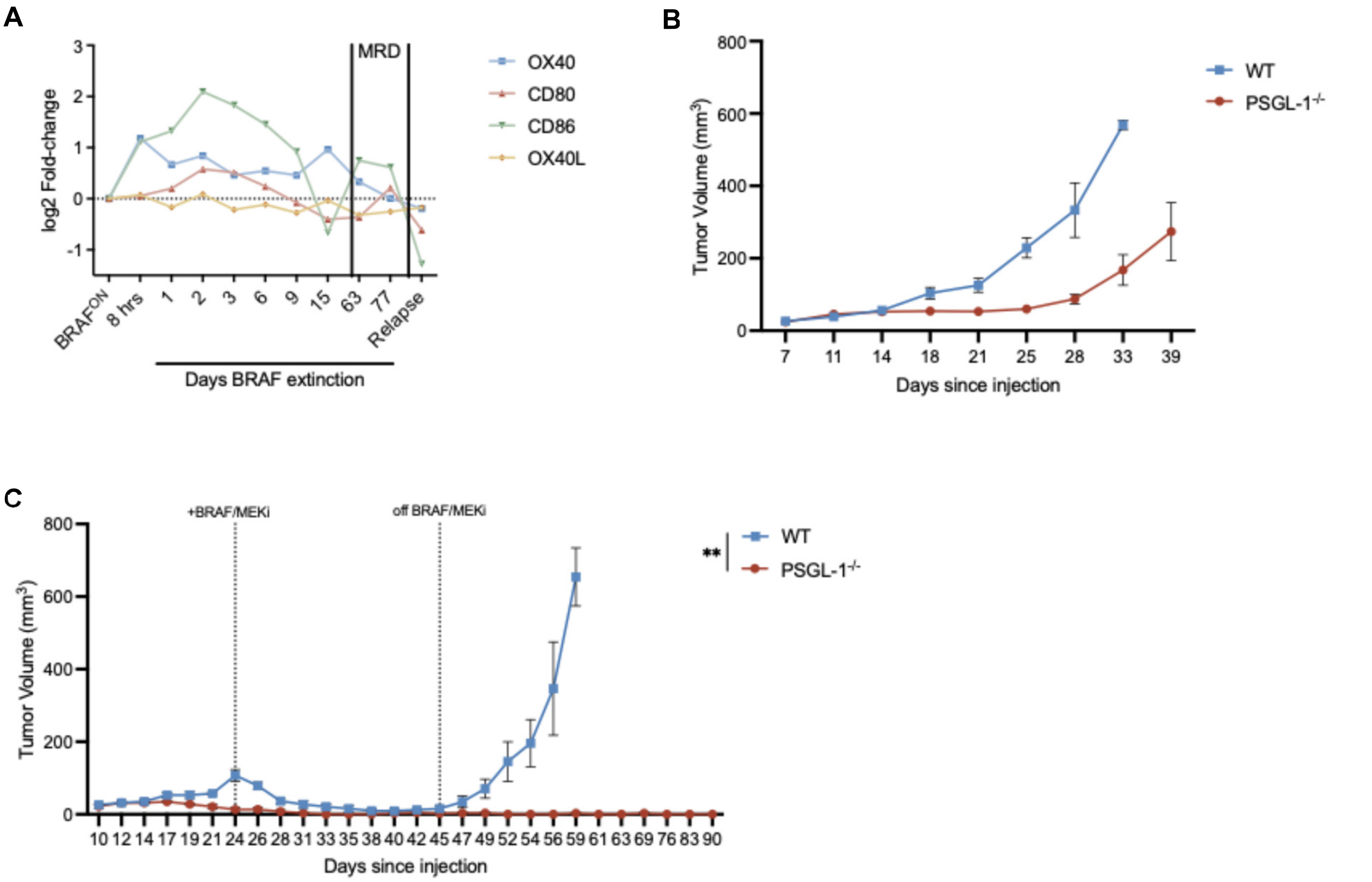


**Supplemental Figure 1. Characterization of MRD and PSGL-1$^{-/-}$ in vivo models. (A)** Remaining 4 out of the top 10 upregulated immune checkpoint after genetic BRAF$^{V600E}$ extinction (days). Data are median centered and reported as log2 fold change. MRD, minimal residual disease. **(B)** Average tumor growth curves of wildtype or PSGL1$^{-/-}$ injected with Yumm1.5 cells (5x10$^{5}$cells) (n=2 or 3 per group). **(C)** Tumor growth curves of wildtype or PSGL1$^{-/-}$ mice +/- BRAF/MEKi Yumm1.5 cells (5x10$^{4}$cells) were intradermally injected in the flank of male mice (n=4 or 5 per group). 24 days after Yumm1.5 injection, BRAF/MEKi or Vehicle control chow were administered. BRAF/MEKi was discontinued after 3 weeks and tumors were monitored for relapse. Dotted line indicates removal of BRAF/MEKi or Vehicle control chow.

### Figure S2

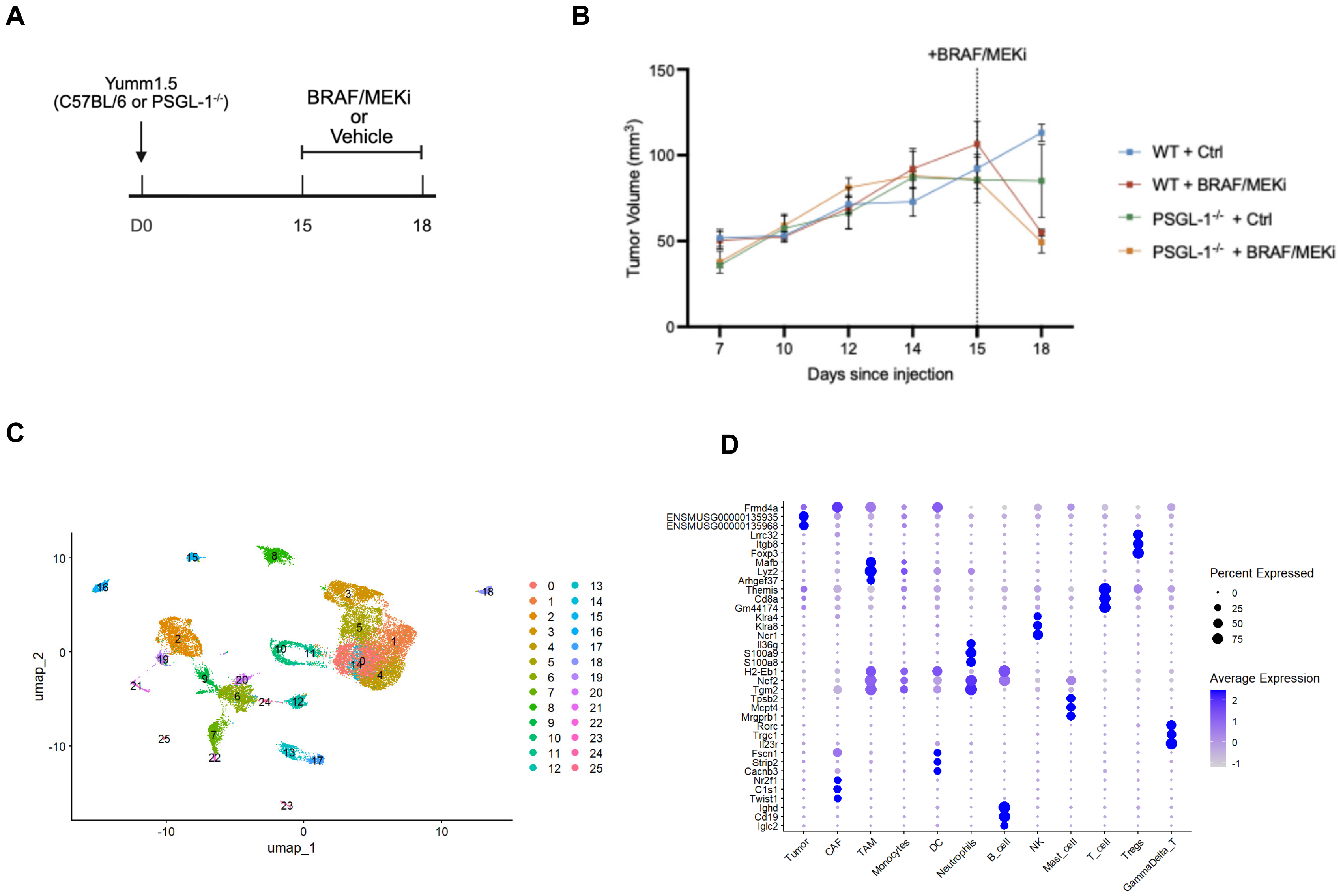


**Supplemental Figure 2. Experimental outline and clustering strategy for scRNASeq analysis. (A)** Experimental timeline of Yumm1.5 tumor-bearing WT or PSGL-1$_{-/-}$ mice treated with either vehicle control or BRAF/MEKi for 72 hours prior to tissue harvest for scRNASeq. Created in https://BioRender.com. **(B)** Tumor growth curves of wildtype (WT) or PSGL1$_{-/-}$mice +/- BRAF/MEKi. Yumm1.5 cells (5x10$_5$cells) were intradermally injected in the flank of male mice (n=3/ group). 15 days after tumor injection, BRAF/MEKi or Vehicle control chow were administered for 72 hours before euthanasia and tissue harvesting for scRNAseq. Data are shown as mean ± SEM. Dotted line indicates administration of BRAF/MEKi or Vehicle control chow. **(C).** UMAP representation of the scRNAseq clusters identified by unsupervised analysis. **(D)** Top maker genes expressed by each assigned major cell subtype.

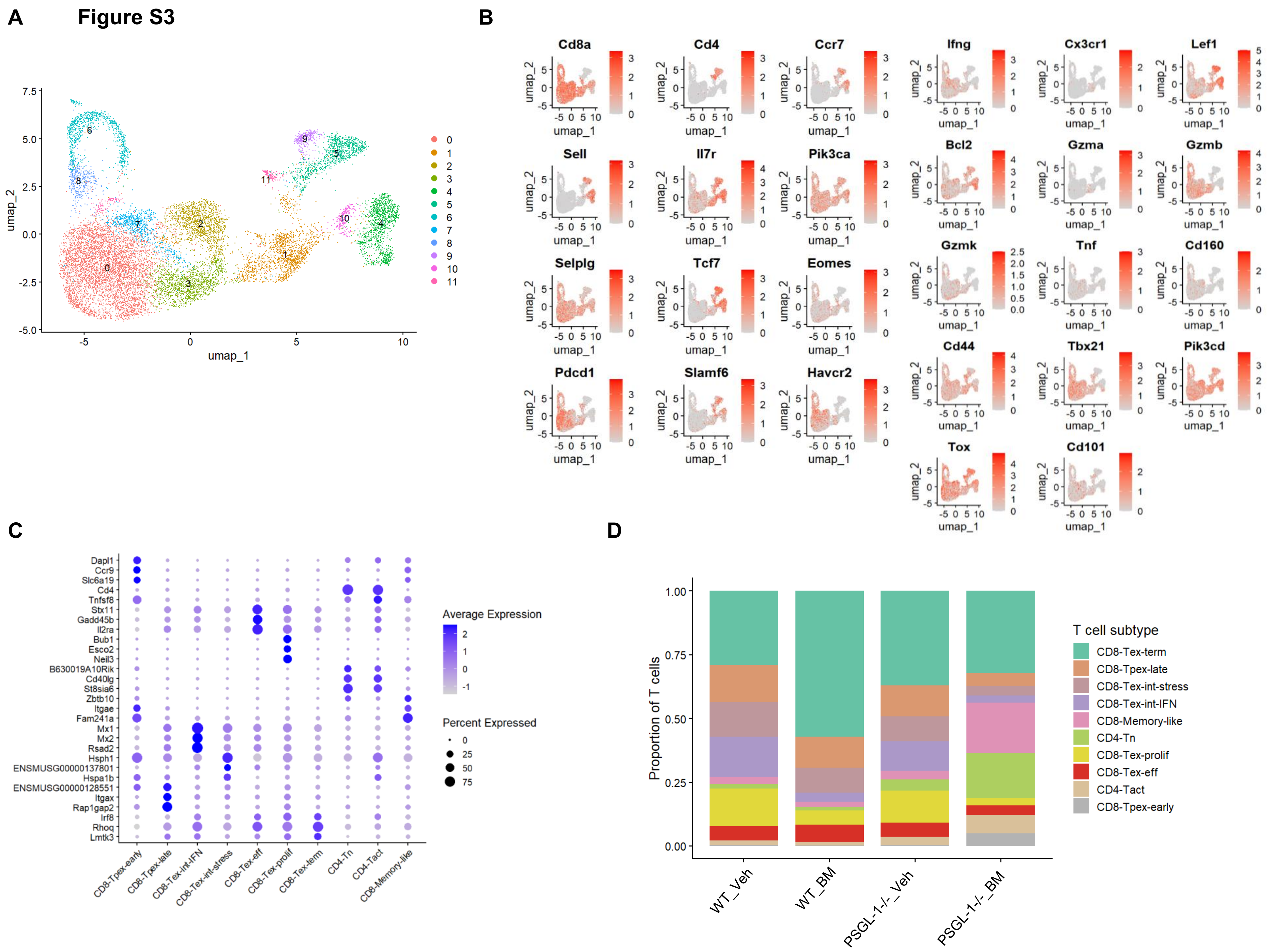


**Supplemental Figure 3. T cell clustering strategy for scRNAseq analysis. (A)** UMAP projection of tumor-infiltrating T cells following reclustering analysis. Unsupervised clustering identified 12 transcriptionally distinct T-cell subpopulations, numbered according to cluster assignment. **(B)** Feature plots showing the expression of representative marker genes used to annotate T-cell subsets, including naïve/memory-associated (*Ccr7, Lef1, Tcf7, Sell*), cytotoxic/effector (*Gzma, Gzmb, Gzmk, Ifng*), progenitor exhausted (*Slamf6, Pdcd1*), terminally exhausted (*Havcr2, Tox, Cd101*), tissue-resident (*Itgae*), and activation-associated markers. Color intensity indicates normalized gene expression.**(C)** Dot plot displaying the top marker genes for each annotated T-cell state. Dot size represents the percentage of cells expressing the indicated gene, and color denotes average normalized expression. **(D)** Relative abundance of annotated T-cell subpopulations across experimental groups (WT_Veh, WT_BM, PSGL-1$^{-/-}$_Veh, and PSGL-1$^{-/-}$_BM). Bars indicate the proportion of total T cells represented by each subset within each condition.

### Figure S4

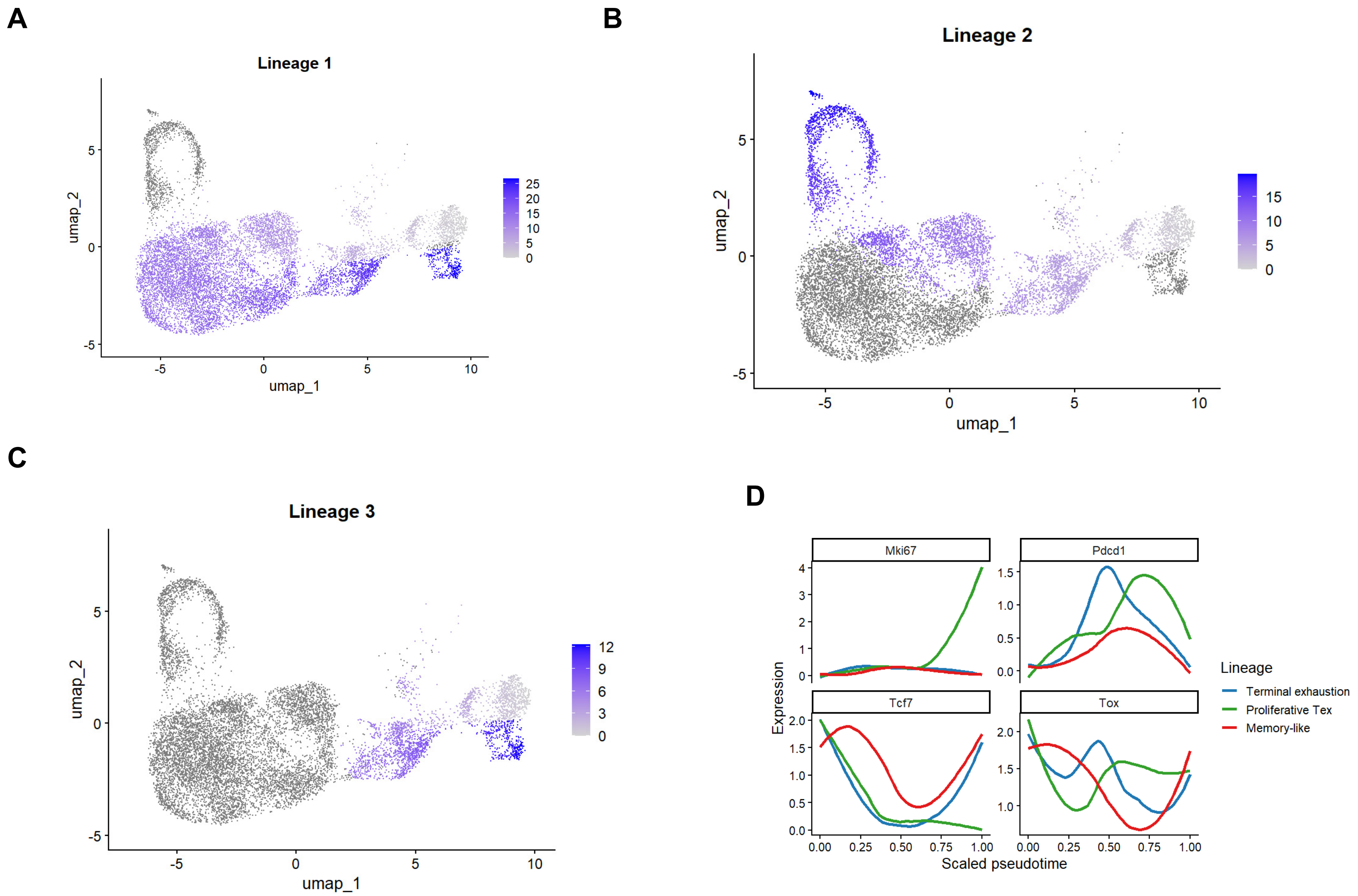


**Supplemental Figure 4. Lineage trajectory analysis of T cell subsets. (A-C)** Visualization of Slingshot-derived pseudotime values projected onto the CD8$^+$ T-cell UMAP for each inferred lineage. Cells assigned to the indicated lineage are colored according to pseudotime progression, with darker blue representing more advanced pseudotime states, while cells not assigned to the lineage are shown in gray. Lineage 1 corresponds to the terminal exhaustion trajectory, Lineage 2 to the proliferative exhausted trajectory, and Lineage 3 to the memory-like trajectory. **(D)** Smoothed expression dynamics of representative lineage-associated genes (*Mki67*, *Pdcd1*, *Tcf7*, and *Tox*) across scaled pseudotime. Curves depict fitted expression trends for terminal exhaustion, proliferative exhausted, and memory-like lineages.

**Figure S5**

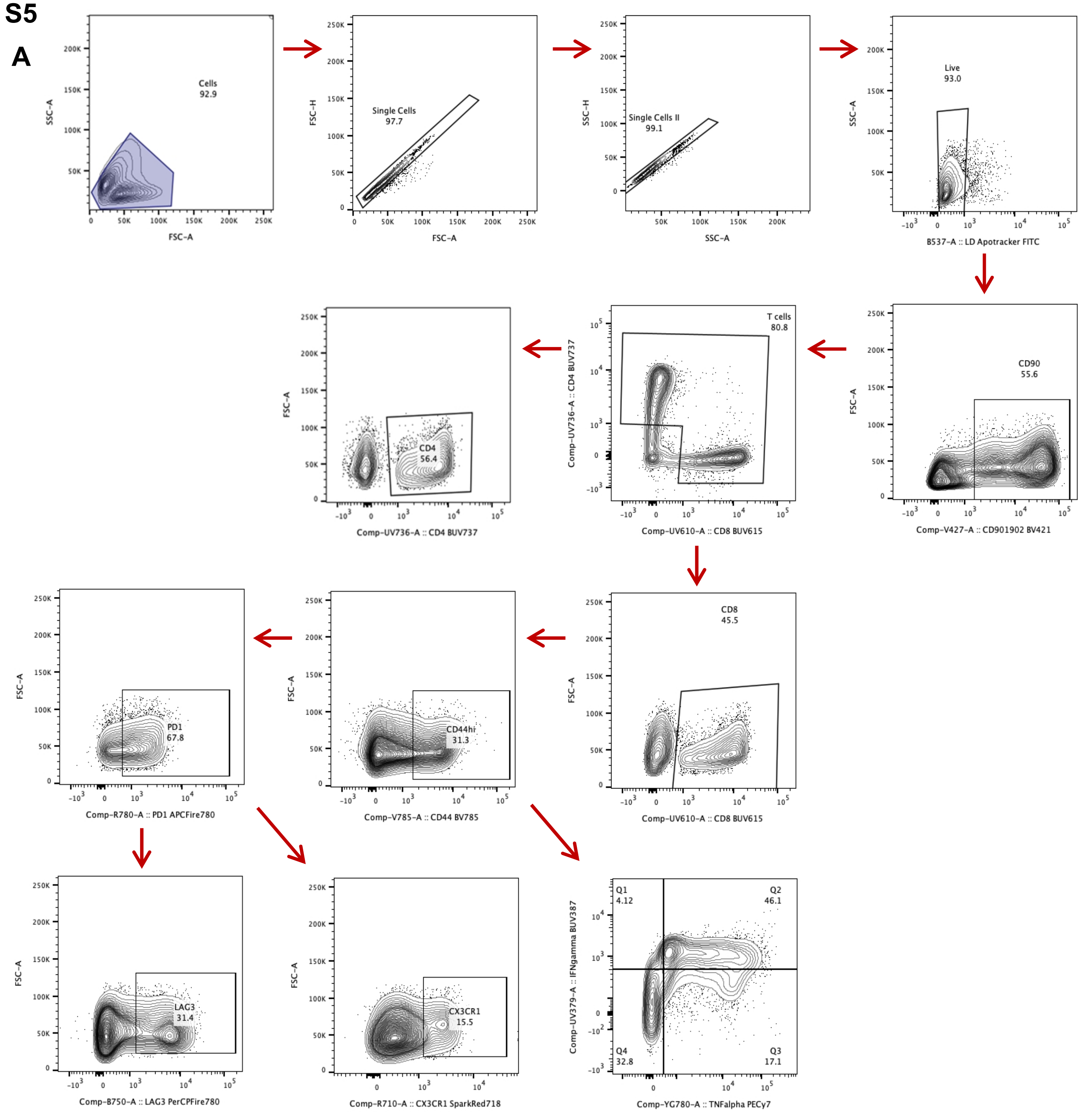


**Supplemental Figure 5. Flow cytometry gating strategy. (A)** Flow cytometry gating strategy used to define CD4⁺ and CD8⁺ T cell populations include activation, exhaustion, and functional markers in CD8⁺ T cells. All analysis was completed using the FlowJo™ software (Version 10.10).

**Figure S6**

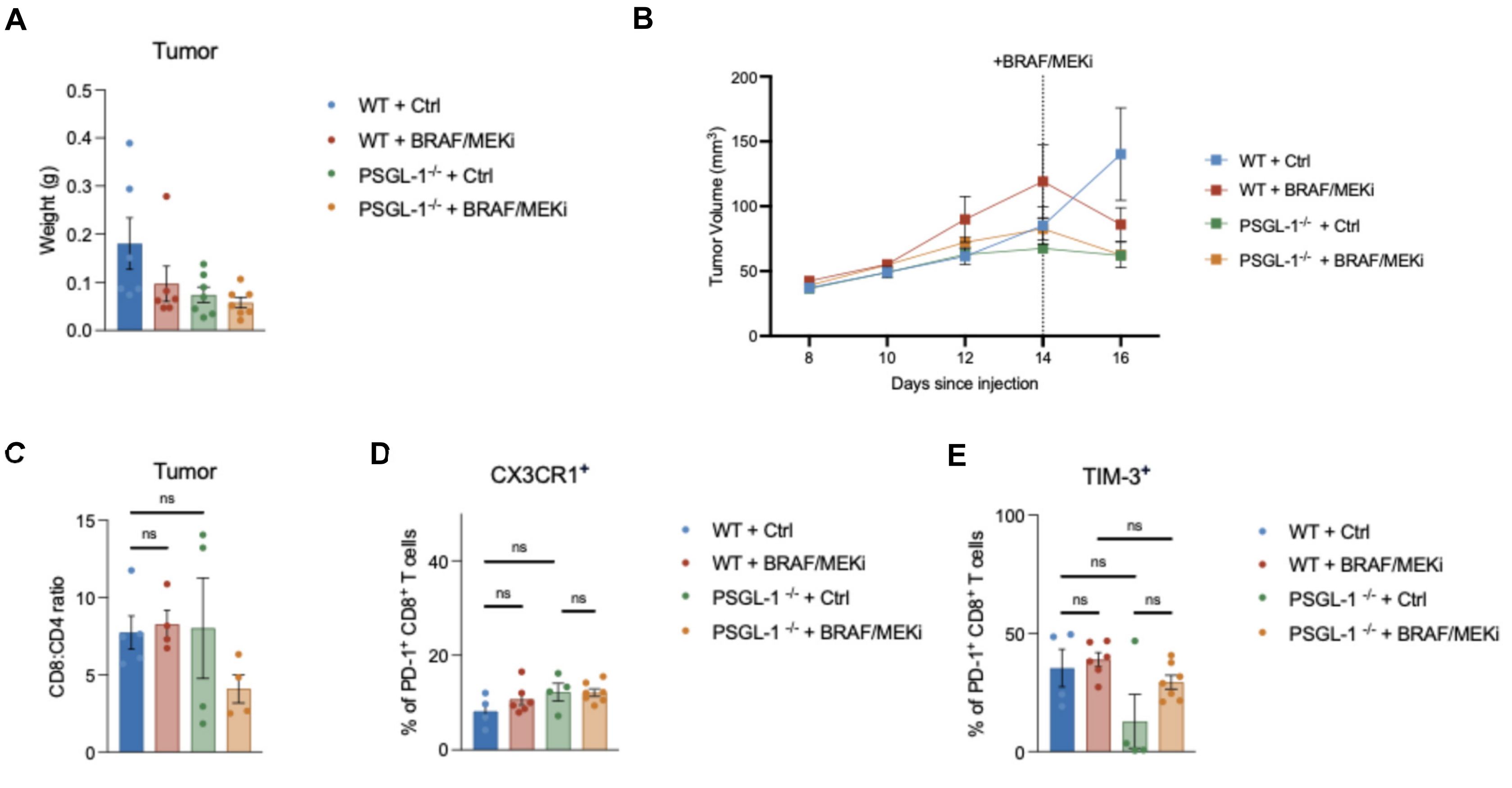


**Supplemental Figure 6. Tumor volume, weight, and tDLN T cell characterization following BRAF/MEK inhibition. (A)** Tumor weight in grams after harvesting from wildtype (WT) or PSGL-1$^{-/-}$ mice treated with Control or BRAF/MEKi chow for 48 hours. **(B)** Tumor growth curves of of wildtype (WT) or PSGL1$^{-/-}$ mice +/- BRAF/MEKi. Yumm1.5 cells ($5x10^5$cells) were intradermally injected in the flank of male mice (n=6 or 7 per group, samples with fewer than $1x10^6$ cells were pooled). 14 days after tumor injection, BRAF/MEKi or Vehicle control chow were administered. Mice were treated with BRAF/MEKi for 48 hours before euthanasia and tissue harvesting. Data are shown as mean ± SEM. Dotted line indicates administration of BRAF/MEKi or Vehicle control chow. **(C)** Ratio of the absolute counts of CD8$^+$ to CD4$^+$ T cells from the tumors of treated mice. CD4$^+$ and CD8$^+$ T cells were gated out of CD90.1$^+$ and CD90.2$^+$ cells from the tumor-draining lymph node of, 48 hours after BRAF/MEKi administration. Cells were stimulated with anti-CD3 and anti-CD28 mAb in the presence of IL-2 and brefeldin A and analyzed via flow cytometry. **(D)** Frequency of CX3CR1$^+$ cells, gated out of PD-1$^+$CD8$^+$ T cells in tdLN following stimulation. **(E)** Frequency of TIM-3-expressing cells, gated out of PD-1$^+$CD8$^+$ T cells in tdLN following stimulation. All data are presented as mean ± SEM. * for $P \leq 0.05$, ** for $P \leq 0.01$, *** for $P \leq 0.001$, and **** for $P \leq 0.0001$, ns (not significant) for $P \geq 0.05$.

### Figure S7

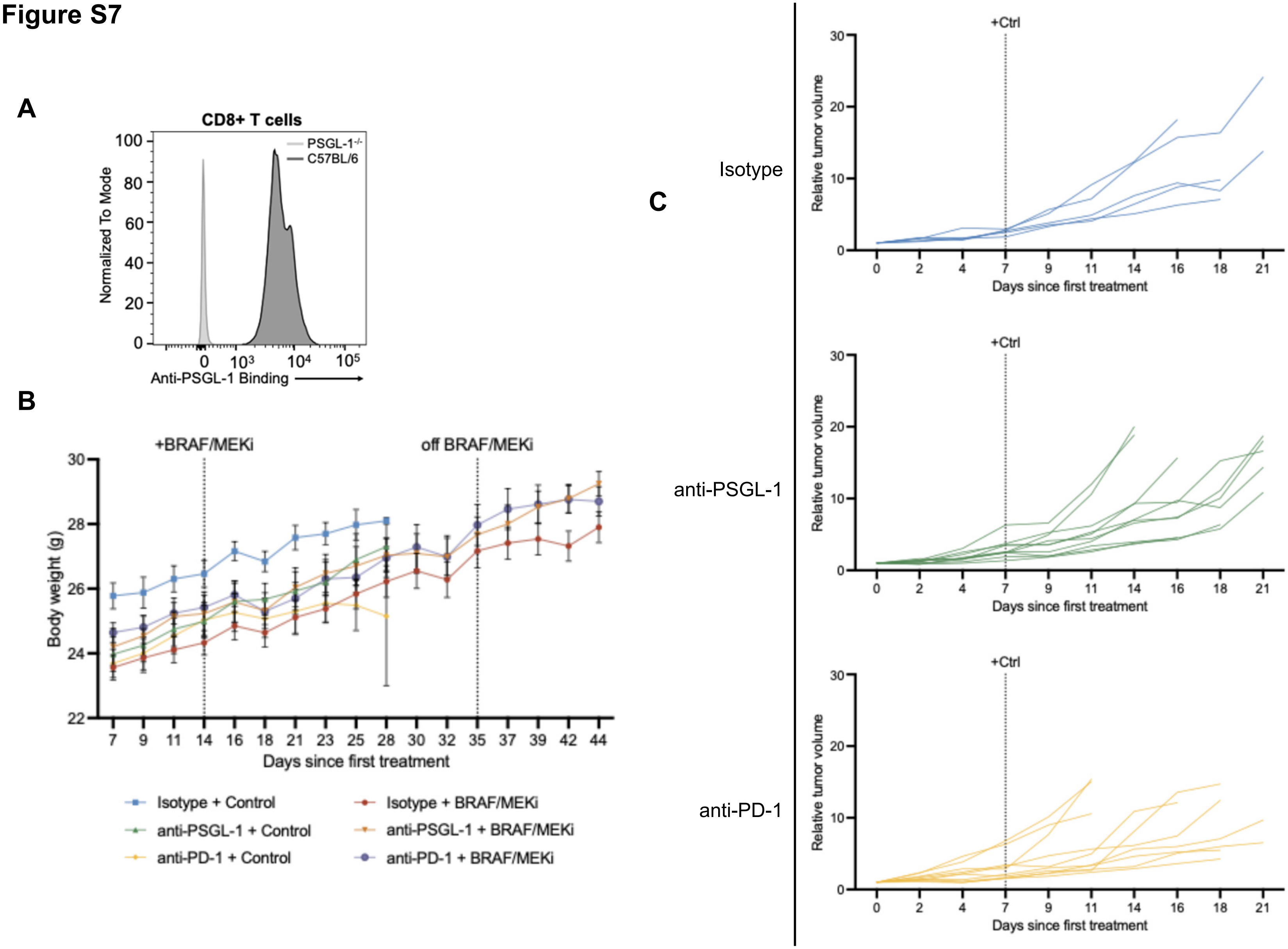


**Supplemental Figure 7. Relative tumor volume of Vehicle-treated mice treated with antibodies targeting PSGL-1 or PD-1 (A)** PSGL-1 specific binding of the anti-mouse PSGL-1 antibody was validated by incubating C57BL/6 or PSGL-1$^{-/-}$ splenocytes with a conjugated anti-mouse IgG secondary antibody for flow cytometric analysis. Shown is anti-PSGL-1 binding out of CD8$^{+}$ T cells. **(B)** Body weights of mice in each group were monitored and measured 3 times per week throughout the duration of the study. **(C)** Relative tumor growth curves of mice treated with either anti-PSGL-1 or anti-PD-1 on vehicle control chow. Tumor volume values were normalized to day 0 on either anti-PSGL-1, anti-PD-1, or Isotype control. Yumm1.5 cells ($5x10^5$cells) were intradermally injected in the flank of male mice (n=5 (Ctrl + Isotype) or 10). Mice were treated with Isotype control, anti-PSGL-1, or anti-PD-1. Tumors were measured and mice weighed 3 times per week. Arrows indicate days that mice were treated with anti-PSGL-1. Isotype or anti-PD-1 treated mice were dosed with respective agent on days 0, 7, 14, and 21. Mice were euthanized at humane endpoint.
